# Robo2 regulates synaptic oxytocin content by affecting actin state

**DOI:** 10.1101/553198

**Authors:** Savani Anbalagan, Janna Blechman, Michael Gliksberg, Ron Rotkopf, Tali Dadosh, Gil Levkowitz

**Affiliations:** Department of Molecular Cell Biology, Weizmann Institute of Science, PO Box 26, Rehovot 7610001, Israel; Bioinformatics Unit, LSCF, Weizmann Institute of Science, PO Box 26, Rehovot 7610001, Israel; Electron Microscopy Unit, Department of Chemical Research Support, Weizmann Institute of Science, PO Box 26, Rehovot 7610001, Israel

**Keywords:** synapse, roundabout, neuropeptide oxytocin, arginine-vasopressin, hypothalamus, neuroendocrine, zebrafish

## Abstract

The regulation of neuropeptide level at the site of release is essential for proper neurophysiological functions. We focused on a prominent neuropeptide, oxytocin (OXT), and used the zebrafish as an *in vivo* model to visualize and quantify OXT content at the resolution of a single synapse. We found that OXT-loaded synapses were surrounded by polymerized actin. Perturbation of actin filaments by cytochalasin-D resulted in decreased synaptic OXT levels. Live imaging of the actin probe, Lifeact-EGFP, showed reduced mobility in OXT synapses in *robo2* mutants, which displayed decreased synaptic OXT content. Using a novel transgenic reporter line allowing real-time monitoring of OXT-loaded vesicles, we showed that *robo2* mutants display slower rate of vesicles accumulation. OXT-specific expression of dominant-negative Cdc42, which is a key regulator of actin dynamics and a downstream effector of Robo2, led to a dose-dependent increase in OXT content in WT, and a dampened effect in *robo2* mutants. Our results link Robo2-Cdc42 signalling, which controls local actin dynamics, with the maintenance of synaptic neuropeptide levels.

## Introduction

The regulation of neurotransmitter level at the site of release is essential for proper neuronal function and requires constant replenishment, capture and removal of excess or aged components in synapses. The axonal cytoskeleton contains microtubules, neurofilaments and actin. Actin dynamics have been studied primarily in the context of morphological changes of growth cones and synapse; however, recent studies indicate that dynamic changes in synaptic actin also modulate synaptic efficacy (Cingolani and Goda, 2008). Conflicting evidence exists for the role of actin filaments in regulating vesicular mobility (Gaffield et al., 2006; Levitan, 2008; Miyamoto, 1995; Nelson et al., 2013; Nunes et al., 2006). However, the majority of such studies were based on actin disrupting or polymerizing agents using cell culture systems or non-vertebrate genetic model organisms. Others have claimed that actin has a molecular scaffolding, not propulsive, role in presynaptic function (Sankaranarayanan et al., 2003).

A well characterized signalling pathway that regulate actin dynamics in neurons is mediated by the Robo family of cell-surface receptors (Kidd et al., 1998; Slovakova et al., 2012). Activation of Robo triggers actin depolymerization which is mediated by intracellular actin interacting regulatory proteins such as Cdc42, N-Wasp and Arp2/3 (Wong et al., 2001).

To address the fundamental topic of synaptic neuropeptide homeostasis we focused on the well-studied neuropeptide, oxytocin (OXT). Oxytocin is a classical evolutionarily conserved neuropeptide, which is involved in the maintenance of various homeostatic functions and whose major axonal release site is the posterior pituitary, also known as the neurohypophysis (Pearson and Placzek, 2013; Wircer et al., 2016). Thus neurosecretory cells produce the neuropeptides vasopressin and oxytocin that are packed into large dense core vesicles (LDCV) and are transported along the axons that terminate in the neurohypophysis. Upon physiological demand, the neuropeptide is released into the blood stream to influence the function of target cells throughout the body (Burbach et al., 2001; Knobloch and Grinevich, 2014). In contrast to small neurotransmitters synaptic release, which mainly occurs in highly specialized membrane structures called active zones, neuropeptides, such as OXT, are release from LDCVs from any part of the neuron, including dendrites and *en passant* axonal synapses (Chini et al., 2017; Leng and Ludwig, 2008; Morris and Pow, 1988). Accordingly, axonal termini of hypothalamic magnocellular OXT neurons converge into the neurohypophysis, where they form numerous *en passant* synapses in a form of highly dense axonal varicosities, also known as axonal swellings or Herring bodies (Tweedle et al., 1989). These structures have been identified as *bona fide* synapses that store OXT-containing LDCV and release them upon physiological demand (Miyata et al., 2001; Wittkowski and Brinkmann, 1974). Thus, in this manuscript, the term “synapses” will henceforth refer to these neurohypophyseal axonal swellings.

Here we used a combination of transgenic zebrafish OXT-specific reporters allowing monitoring and quantification of synaptic OXT levels. We associate between synaptic actin state and OXT accumulation at the site of release. We further show that Robo2-Cdc42 signaling, which has previously been associated with modulation of actin polymerization in the growth cones of guided axons, regulates synaptic actin state and OXT neuropeptide accumulation in the neurohypophyseal termini.

## Results

### Quantitative analysis of synaptic OXT neuropeptide levels *in vivo*

The optically transparent zebrafish larva has a few dozens of OXT neurons, which enables analyzing the function of each neuron down to the single-synapse resolution in the context of a living vertebrate animal (Blechman et al., 2011; Gutnick et al., 2011). To visualize and quantify synaptic OXT neuropeptide content at the resolution of a single synapse, we combined anti-OXT antibody staining with a transgenic reporter zebrafish, *Tg(oxt:EGFP)*, in which neurohypophyseal OXT synapses are filled with cytoplasmic EGFP (**Fig. 1A**). In this manner, the structure of the synapse itself, labelled by EGFP, could be differentiated from it’s content of oxytocinergic LDCVs, labelled by the anti-OXT antibody. We used image thresholding settings that allowed detection of individual EGFP-labelled synapses and their neuropeptide content, which appeared in the form of immune-reactive OXT puncta that colocalized with these EGFP-labelled synapses (**Fig. 1A and Figure 1—video supplement 1**).

**Figure 1 with 1 supplement.**
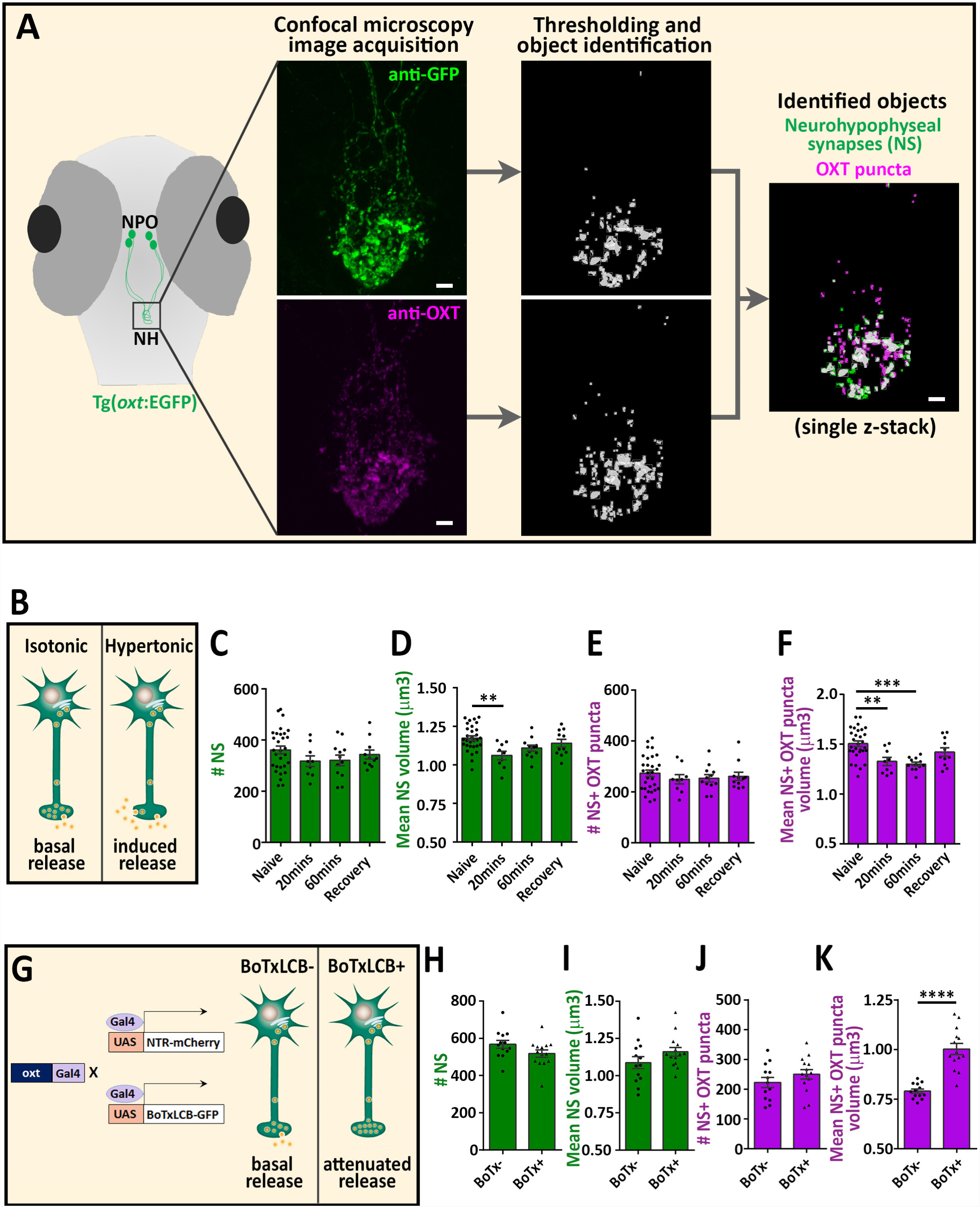
*In vivo* quantification of synaptic OXT content in neurohypophyseal axonal termini. **(A)** Scheme describing the pipeline for detection of neurohypophyseal synapses (NS). Whole-mount imaging of hypophysis of 5 days post-fertilization (dpf) transgenic reporter Tg(*oxt*:EGFP) zebrafish following immunostaining with anti-EGFP and a specific antibody against endogenous OXT protein. Analysis of GFP-positive neurohypophyseal synapses (NS) and OXT puncta were performed by using the “object identifier” function in Volocity software on individual channels. **(B-F)** Hypertonic challenge induces synaptic OXT release. Transgenic Tg(*oxt*:EGFP) larvae at 8 dpf were treated with hypertonic solution (25% artificial sea salt in Danieau buffer) for 20 or 60 min. The larvae that underwent 60 min treatment were allowed to recover in isotonic Danieau buffer for another 60 min. The mean number and volume of NS (**C,D**) and OXT puncta overlapping with GFP+ NS (**E,F**) were quantified (**P<0.01 and ***P<0.001; one-way ANOVA). **(G-K)** Cell-specific blockage of synaptic release using botulinum toxin light chain B increases hypophyseal OXT levels (**G**). The mean number and volume of mCherry+NS (**H,I**) and oxytocin puncta overlapping with mcherry+ NS (**J,K**) were quantified in Tg(*oxt*:Gal4; UAS:NTRmCherry) labelled as BoTx(n=13 larvae) versus Tg(*oxt*:Gal4; UAS:BoTxLCB-GFP; UAS:NTRmCherry) labelled as BoTx+ (n=14 larvae) larvae at 8 dpf (****P<0.0001; Student’s *t*-test). Error bars indicate SEM in (C-F, H-K).

To validate our detection method, we subjected the fish to hypertonic osmotic challenge (25% sea salt) (**Fig. 1B**), which is known to induce the release of oxytocin and vasopressin and, consequently reduced neuropeptide content in the pituitary of both mammals and fish (Balment et al., 1980; Huang et al., 1996; Leng and Russell, 2018; Pierson et al., 1995). We detected several hundreds of synapses and OXT-stained puncta in the neurohypophysis of naïve 8 days post-fertilization (dpf) zebrafish larvae. While the number of these EGFP-positive synapses remained unaltered following acute hypertonic challenge, synaptic and OXT puncta volumes were decreased (**Fig. 1C-F**). Recovery of larvae from hypertonic to isotonic condition (for 1 hour) led to reversal of the observed phenotypes, namely increase in size of the synapses and OXT puncta (**Fig. 1D and F**). These results are in agreement with the reported hypertonicity-induced cell shrinking mechanisms essential for osmosensation in these neurons (Prager-Khoutorsky et al., 2014).

To further validate our method, we demonstrated that inhibition of tonic OXT release by oxytocin neurons-specific expression of botulinum toxin, which cleave the vesicle-associated membrane protein (VAMP) thereby inhibiting LDCVs synaptic release. We utilized a conditional triple transgenic line *Tg*(*oxt:Gal4; UAS:NTR-mCherry; UAS:BoTxLCC-GFP*), in which synaptic vesicle release from OXT neurons was blocked due to intracellular expression of botulinum toxin light chain B (BoTxLCB) fused to GFP (Sternberg et al., 2016), and the synapses were labelled via mCherry (**Fig.1G**). OXT-specific expression of BoTxLCB lead to increased volume of OXT puncta, but without affecting the number and volume of the synapses, indicative of accumulation of OXT content (**Fig. 1H-K**). We conclude that the above methodology can faithfully quantify synaptic neuropeptide homeostasis *in vivo*.

### Disruption of F-actin affects axonal swellings size and OXT neuropeptide content

To study the spatial relationship between actin and synaptic OXT neuropeptide we performed super-resolution microscopy of a double transgenic line, *Tg(oxt:Gal4; UAS:Lifeact-EGFP)*, in which the filamentous actin (F-actin) probe, Lifeact-EGFP, was specifically expressed in OXT neurons. *Tg(oxt:Gal4; UAS:Lifeact-EGFP)* larvae were immunostained with antibodies against EGFP and endogenous OXT and imaged using stochastic optical reconstruction microscopy (STORM) (**Fig. 2A-D**). Imaging revealed various characteristics of actin and neuropeptides. Lifeact-EGFP signal exhibited a cage-like structure engulfing OXT in neurohypophyseal synapses (**Fig. 2B,C and Fig. 2 - supplemental video 1**). In some synapses, actin-associated neuropeptides were observed only in the synaptic periphery, suggesting that the actin-coated vesicles are associated with the plasma membrane of such synapses (**Fig. 2D**). These results are in agreement with previous reports that actin filaments form a cage-like structure associated with synaptic vesicles thus restricting the mobility of these vesicles (Hirokawa et al., 1989; Miyamoto, 1995).

**Figure 2 with 1 supplement.**
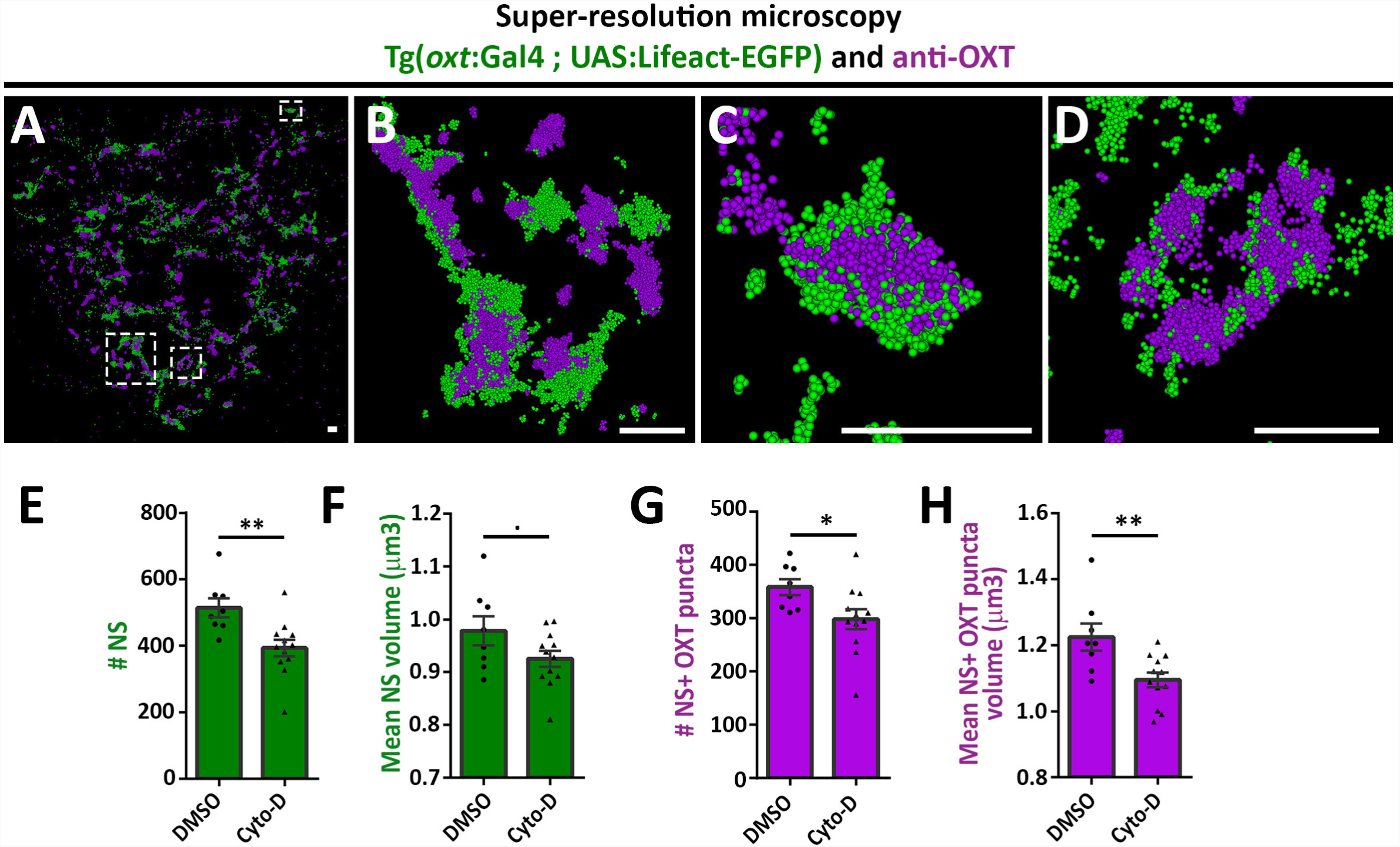
Actin is required for synaptic OXT homeostasis. (**A-D**) Spatial relationship between actin and neuropeptide in hypophysis revealed by super-resolution microscopy. Stochastic optical reconstruction microscopy (STORM) images of the neurohypophyseal area of 6-days post-fertilization (dpf) Tg(*oxt*:gal4; UAS:lifeact-EGFP) larvae stained using anti-OXT and anti-GFP antibodies (**A**). **B-D:** Magnifications of squared areas. Scale bars: 1 µm. (**E-H**) Assessment of the effect of cytochalasin D treatment on synaptic properties in Tg(*oxt*:EGFP) larvae. Larvae were treated with DMSO or cytochalasin D at 400 nM between 4 and 5 dpf, stained using anti-GFP and anti-OXT antibodies and quantified following imaging as in Figure 1. Treatment with cytochalasin D led to decreased number of neurohypophyseal synapses (NS) (**E**) and associated OXT puncta (**G**) and decreased mean volume of NS (**F**) and associated OXT puncta (**H**) (E-H; P<0.01; Student’s *t*-test; n= 8, 12 for DMSO and cytochalasin D treatment respectively). Error bars indicate SEM in (E-H).

To investigate the role of actin in capture of oxytocin vesicles in hypophyseal synapses, we perturbed the hypophyseal synaptic actin filaments by using the actin depolymerizing agent cytochalasin D (Cooper, 1987). Transgenic Tg(*oxt*:EGFP) larvae were treated with cytochalasin D between 4 and 5 days post-fertilization at a concentration of 400nM, which is a tolerated dose that allows normal development and viability of zebrafish larvae (Trendowski et al., 2014). Subsequent analysis revealed that the number and size of the synaptic axonal swellings was reduced upon cytochalasin D treatment (**Fig. 2E,F**), which is in agreement with the involvement of F-actin in regulating the synaptic morphology (Zhang and Benson, 2001). Cytochalasin D treatment also affected both the number and volume of OXT puncta (**Fig. 2G,H**). These results suggest that actin polymerization is required for axonal swelling morphogenesis as well as for homeostasis of synaptic OXT neuropeptide content.

### Robo2 regulates synaptic actin state and OXT accumulation

Next, we searched for candidate signalling pathways that could regulate axonal F-actin in OXT neurons. Robo2 is localized to axonal growth cones and is known to regulate axonal guidance by modulating actin dynamics via other actin interacting regulatory proteins (Kidd et al., 1998; Slovakova et al., 2012). Fluorescent in situ hybridization of *robo2* mRNA showed that Robo2 is expressed in OXT neurons (**Fig. 3A**). To investigate if Robo2 regulates synaptic actin state, we used the zebrafish *robo2-*deficient mutant, *astray* [*robo2*^272z/272z^ (Fricke et al., 2001)], to perform fluorescence recovering after photobleaching (FRAP) on individual OXT synapses expressing the F-actin sensor *Tg(oxt:Gal4;UAS:Lifeact-EGFP)* in either *robo2+/+* or *robo2-/-* animals (**Fig.3B,C**). We reasoned that the dynamics of Lifeact-EGFP fluorescence recovery indicates changes in synaptic polymerized actin, which is available for Lifeact-EGFP binding. Thus, it is expected that synapses wherein actin filaments are highly stable would display an increased time of Lifeact-EGFP fluorescence recovery. Indeed, in comparison to the *robo2*+/+ zebrafish, the recovery of Lifeact-EGFP fluorescence was attenuated in OXT synapses of *robo2*-/-mutants (**Fig. 3D**), which exhibited decreased dynamic Lifeact-EGFP fraction and increased stable fraction (**Fig. 3E,F**). This suggests that Robo2 signalling regulates synaptic actin state of neurohypophyseal OXT neurons.

**Figure 3.**
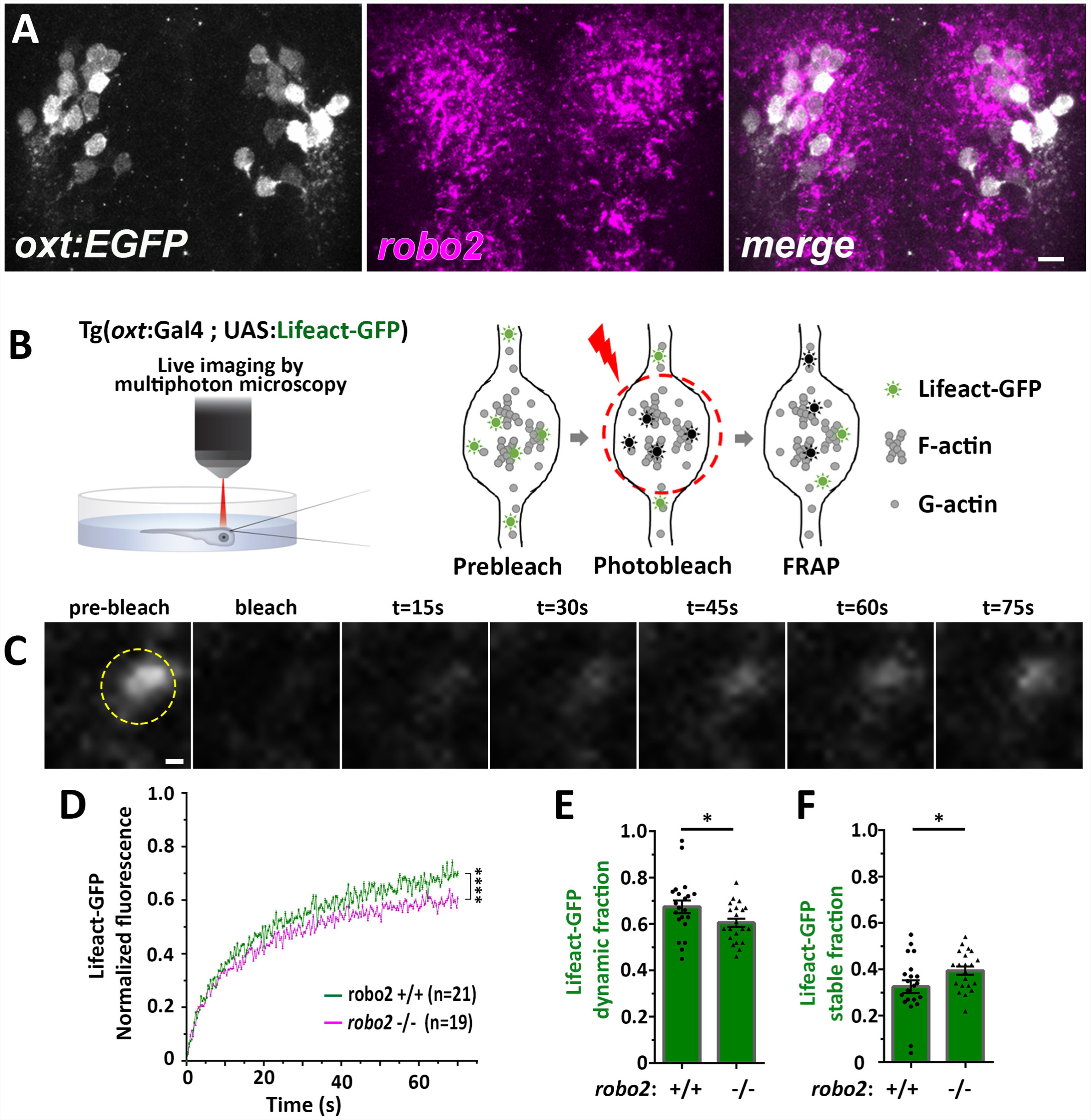
Robo2 is required to regulate synaptic actin states. (**A**) *robo2* is expressed in larval zebrafish neurosecretory preoptic area (NPO) and colocalizes with Oxytocin neurons. Confocal Z-stack images showing fluorescent *in situ* hybridization (FISH) of transgenic larvae Tg(*oxt*:EGFP) (3 days post-fertilization (dpf)) using probes directed against *robo2* mRNAs (magenta), followed by anti-EGFP staining. The NPO area in which OXT neurons were labelled is shown. Scale bar: 20 µm. (**B,C**) Real-time monitoring of synaptic actin dynamics in live transgenic reporter Tg(*oxt*:Gal4; UAS:Lifeact-EGFP) larvae mounted in 0.1% low-melt agarose and imaged using multi-photon microscopy upon Fluorescence Recovery after Photobleaching (FRAP) (**B**). Time-series images of FRAP experiment in a neurohypophyseal synapse with Lifeact-EGFP expression (**C**). Scale bar: 200 nm. (**D-F**) Assessment of synaptic actin dynamics in *robo2* mutant using the transgenic actin dynamics reporter Tg(*oxt*:Gal4; UAS:Lifeact-EGFP) larvae. Graph showing the normalized FRAP profile of Lifeact-EGFP fluorescence intensity in 6-dpf *robo2*^+/+^ (n=21 synapses) and *robo2*^-/^(n=19 synapses) larvae (**D**) (****P<0.0001; two-way ANOVA). Bar graphs showing the dynamic (**E**) and stable (**F**) Lifeact-EGFP fractions in *robo2*^+/+^ vs *robo2*^-/-^neurohypophyseal synapses (*P<0.05; Student’s *t*-test). Error bars indicate SEM in (D-F).

We next asked if Robo2 plays a role in synaptic accumulation of OXT by examining whether neurohypophyseal neuropeptide content is altered in the *robo2-*defiecient mutant zebrafish. We found that the volume of OXT puncta was smaller in *robo2*-/- fish compared to WT controls, while the number and size of axonal swellings were largely unaffected, suggesting that Robo2 is required to maintain synaptic oxytocin levels without affecting synaptic morphogenesis (**Fig. 4A-D**). This phenotype was not due to OXT axonal guidance deficits, as the number of neurohypophyseal OXT axonal projections was similar between *robo2-/-* and *robo2+/+* fish larvae (**Fig. 4E**).

**Figure 4 with 2 supplements.**
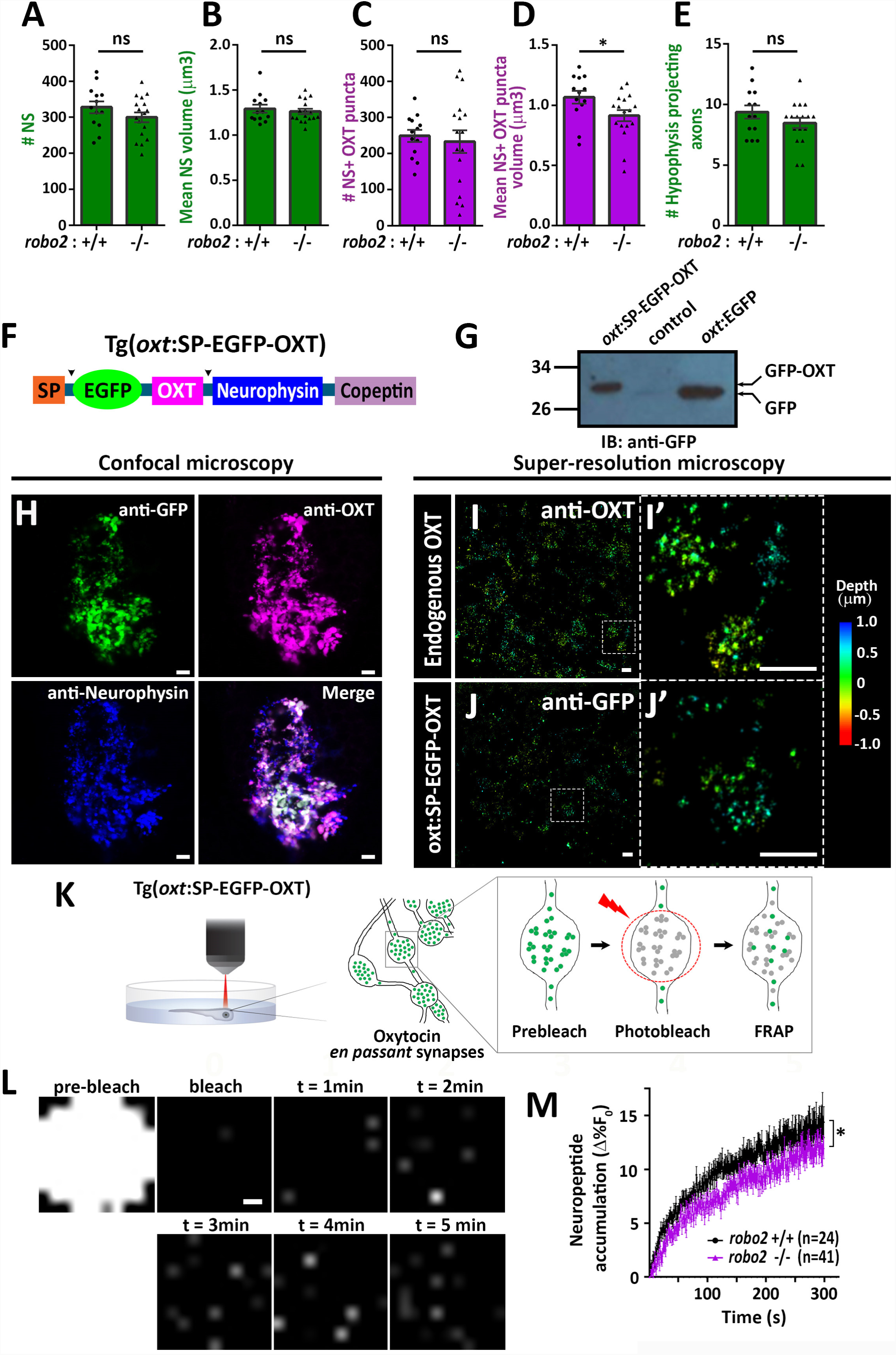
Robo2 is required to regulate synaptic OXT levels. (**A-E**) Assessment of synaptic oxytocin content in *robo2* mutant was performed as described in Figure 1. Graph showing the number and size of neurohypophyseal synapses (NS) (**A,B**) and colocalizing OXT puncta (**C,D**) and the number of neurohypophyseal projecting axons (**E**) in 8-days post-fertilization (dpf) *robo2*+/+ (n=13) vs *robo2*-/-(n=17) larvae (* p<0.05; ns denotes not significant, Student’s *t*-test). **(F)** Schematic of novel transgenic OXT tool Tg(*oxt*:OXT-SP-EGFP-OXT), in which *oxt* promoter drives expression of Oxytocin precursor protein with an internally-tagged EGFP at the C-terminus of the signal peptide. **(G)** Validation of OXT-fusion EGFP protein expression by Western blot analysis. Pituitary protein extracts from adult Tg(*oxt*:OXT-SP-EGFP-OXT), TL (control) and Tg(*oxt*:EGFP) zebrafish were immunoblotted using anti-GFP. **(H)** Validation of OXT-fusion EGFP protein expression by immunohistochemistry. Confocal Z-stack images of the neurohypophyseal area of 5-dpf transgenic Tg(*oxt*:OXT-SP-EGFP-OXT) larvae. Larvae were stained using anti-GFP, anti-OXT and anti-neurophysin antibodies. Scale bar: 1 µm. (**I-J**) Assessment of endogenous vs OXT-fusion EGFP protein localization and packaging by super-resolution microscopy. STORM Z-stack images of the endogenous synaptic OXT (I) in comparison to OXT-EGFP fusion in transgenic Tg(*oxt*:OXT-SP-EGFP-OXT) larvae (**J**). 5 dpf larvae were stained with either anti-OXT (**I**) or anti-GFP (**J**) antibodies and visualized by STORM as described in the Method section. Scale bar:. The right panels I’ and J’ show magnifications of the represented areas outlined in **I** and **J.** Scale bars: 1 µm 200 nm (I’ and J’). Colour code indicates Z-axis depth. (**K,L**) Real-time monitoring of synaptic oxytocin vesicle dynamics in live transgenic reporter Tg(*oxt*:OXT-SP-EGFP-OXT) larvae. Schemata of the experimental design (**K**): 6 dpf larvae were mounted dorsally in low-melt agarose gel, submerged in E3 embryo buffer and imaged using multi-photon microscopy. Neurohypophyseal synapses containing OXT-EGFP vesicles were photobleached and the fluorescence recovery after photo bleaching (FRAP) over time was monitored. The fluorescent recovery occurs due to the dynamic exchange of mobile and unbleached OXT-EGFP vesicles from neighbouring *en passant* synapses over time (t=1-5 min). The brightness and contrast of the images were increased to visualize the individual pixels (**L**). Scale bar: 200 nm. (**M**) Graph showing neuropeptide accumulation (normalized FRAP curves) of OXT-EGFP in 6-dpf *robo2*+/+ (n=24) vs robo2-/-(n=41) neurohypophyseal synapses (*P<0.05, two-way ANOVA). Neuropeptide accumulation represents full scale normalized data to account for differences in synaptic OXT-EGFP fluorescence, i.e. fluorescent values upon photobleaching were normalized to zero. Error bars indicate SEM in (**A-E** and **M**).

**Figure 4 - figure supplement 1.**
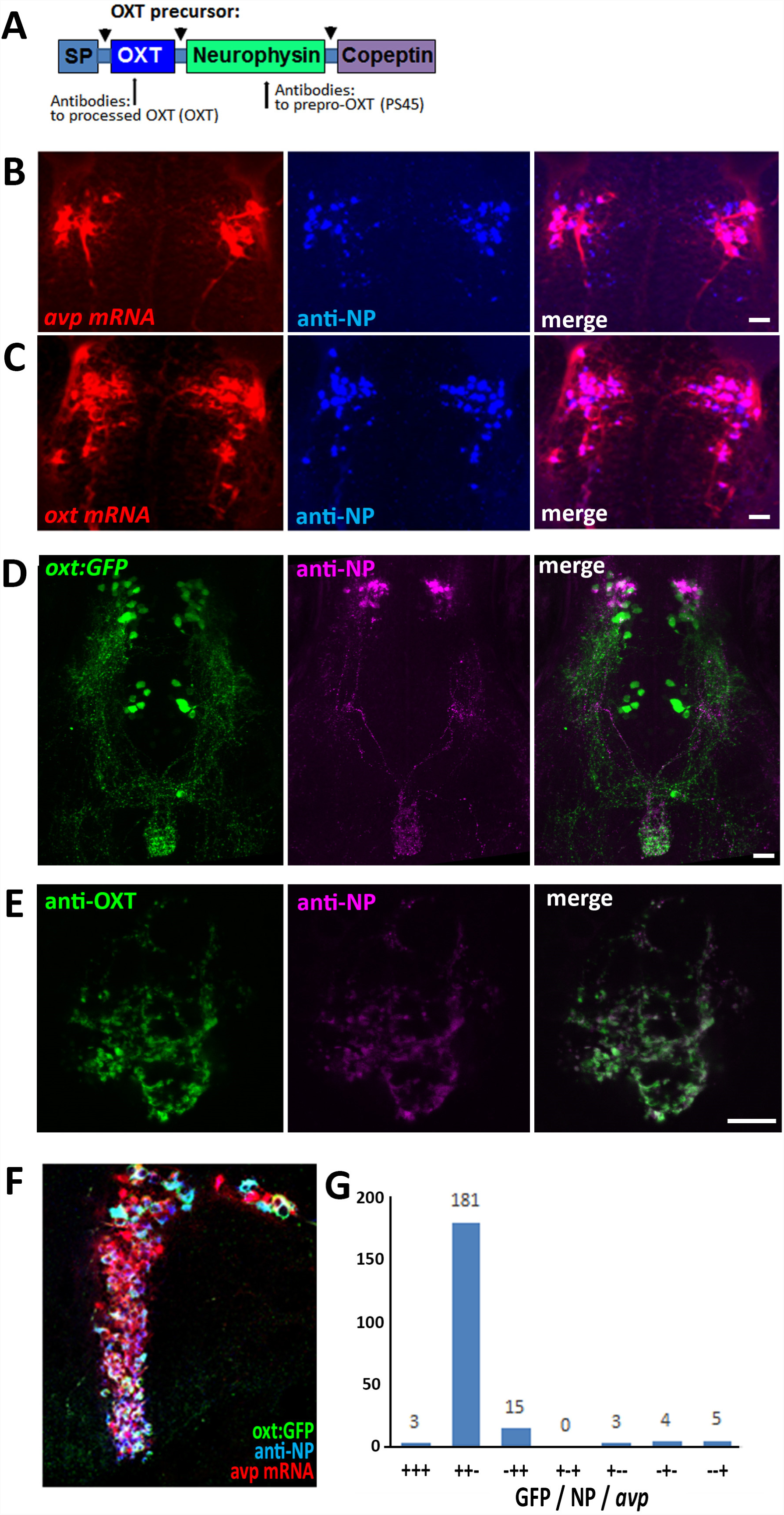
(**A**) Schema of protein coded by *oxt* gene. Antibodies anti-OXT and PS45 were targeted to processed OXT and neurophysin respectively. SP denotes signal peptide, arrow indicates cleavage site. Validation of the EGFP-OXT fusion protein is shown in **Fig. 3H**. (**B-G**) Confocal Z-stack images showing Fluorescenct in situ hybdridization (FISH) of 3-days post-fertilization (dpf) larvae using probes for *avp* (**B**) or *oxt* (**C**) mRNA (red), followed by PS45 staining. The NPO area in which oxytocin neurons were labelled is shown. Confocal maximum intensity projection (MIP) images showing immunostaining of 5-dpf old transgenic Tg(*oxt*:EGFP) larvae using antibodies directed against GFP and neurophysin (NP). The whole neurohypophyseal tract (**D**) and the hypophyseal projections (**E**) are shown. Confocal MIP images showing FISH and immunostaining of 3-months-old adult transgenic Tg(*oxt*:EGFP) zebrafish using probes directed against *avp* and immunostaining against EGFP and neurophysin (NP). Quantification of number of cells positive for GFP or PS45 or *avp* is shown (**G**).

To visualize OXT neuropeptide dynamics in real time, we developed a novel transgenic tool, *Tg[oxt:OXTSP-EGFP-OXT-NP]*, in which OXT promoter drives the expression of EGFP fused with the OXT precursor, between the signal sequence and the OXT peptide (**Fig. 4F**). Efficient generation of a cleaved EGFP-OXT fusion protein was confirmed by Western blot analysis of pituitary synaptosomes isolated from adult pituitaries of transgenic fish (**Fig. 4G**). The vesicular synaptic expression of the EGFP-OXT fusion protein was validated by triple coimmunostaining of the transgenic EGFP-tagged product, together with two specific antibodies to the endogenous neurophysin and OXT, which revealed that the EGFP-tagged OXT transgene mainly co-localized with the mature (i.e. cleaved and cyclised) OXT neuropeptide (**Fig. 4H** and (**Fig.4 – Fig. supplement 1**)). As expected, in some cases, the OXT-GFP reporter co-localized with vesicles containing both cleaved and OXT-neurophysin protein, suggesting that it reported an intermediate step in the process (**Fig. 4H**). To verify the vesicular synaptic localization of the EGFP-tagged OXT we performed super-resolution microscopy (STORM) imaging of *Tg[oxt:OXTSP-EGFP-OXT-NP]* larvae. Our results showed that similar to the endogenous OXT, the EGFP-OXT reporter exhibited a clustered organization, indicative of large dense core vesicular organization of OXT (**Fig. 4I,J** and **Supplement video 1**).

To assess the vesicular mobility and monitor *in vivo* neuropeptide homeostasis using our novel transgenic tool, we performed FRAP analysis of individual neurohypophyseal swellings using two-photon microscopy in live 6- days post-fertilization transgenic *Tg[oxt:OXTSP-EGFPOXT-NP]* larvae (**Fig. 4K**). Upon bleaching, we observed gradual recovery of OXT-EGFP fluorescence indicating the mobilization of transiting OXT-loaded vesicles in the synapses. The extent of fluorescence recovery was low (13%), suggesting that the majority of the bleached OXT-EGFP-positive vesicles were stationary and not mobile (**Fig. 4L)**. FRAP analysis in transgenic larvae on the background of *robo2* mutants revealed that the fluorescence recovery rate in *robo2*-/- mutants, was significantly lower than in *robo2*+/+ larvae (**Fig. 4M**). Taken together, these results suggest that Robo2-mediated signaling regulates actin dynamics as well as the accumulation of oxytocin-containing vesicles in neurohypophyseal synapses.

### Robo2 and Cdc42 regulate OXT neuropeptide levels

We hypothesized that Robo2 exerts its effect on OXT content via actin polymerization; thus we search for a candidate signalling mediator that could link between these two Robo2-mediated processes. Robo signalling is known to affect the transition of the Rho-GTPase protein Cdc42 from GTP- to GDP- bound state, resulting in decreased actin polymerization (Wong et al., 2001). We therefore tested if conditional OXT-specific expression of dominant-negative (i.e. GDP-bound) mutant form of CDC42, termed Cdc42(T17N), would affect synaptic OXT content and whether the effect would be dampened in a Robo2 mutant. We used our Tg(*oxt*:Gal4) transgenic driver to drive specific oxytocinergic expression of EGFP-Cdc42(T17N) fusion protein, which was regulated by ten Gal4 DNA binding UAS repeats (Ando et al., 2013). Thus, one cell-stage embryos were co-injected with transposon-based DNA constructs harboring *oxt*:Gal4 together with either UAS:EGFP-Cdc42-T17N or UAS:EGFP construct as a control. (**Fig. 5A**). We then quantified OXT content and EGFP-Cdc42(T17N) protein in each synapse (**Fig. 5B**). We took advantage of the variable expression of the injected construct in each individual synapse, to examine whether differences in OXT content correlated with expression levels of EGFP-Cdc42(T17N) or the control EGFP. Regression analysis of OXT fluorescence as a function of EGFP fluorescence in each injected zebrafish larvae showed that high EGFP fluorescence led to decreased OXT levels in both *robo2+/+* and *robo2-/-* larvae (**Fig. 5C;** p<0.01, adj. R^2^=0.93 and 0.85 respectively) (**Fig. 5C**). This effect is likely due to overexpression of the untethered EGFP. In contrast, expression levels of EGFP-Cdc42(T17N) are positively correlated with increased OXT content in *robo2+/+* (**Fig. 5D**; p<0.01, adj. R^2^=0.74). However, the positive effect of EGFP-Cdc42(T17N) on synaptic OXT content was dampened in *robo2-/-* mutants (**Fig. 5D**; p=0.35, Adj. R^2^=0.52), a finding which is consistent with the fact that Robo2 inactivation leads to increased levels of active GTP-bound Cdc42. These results place the small GTPase Cdc42, which is a key regulator of actin dynamics, in a Robo2 signalling cascade that controls the levels synaptic OXT neuropeptide (**Fig. 5E**).

**Figure 5.**
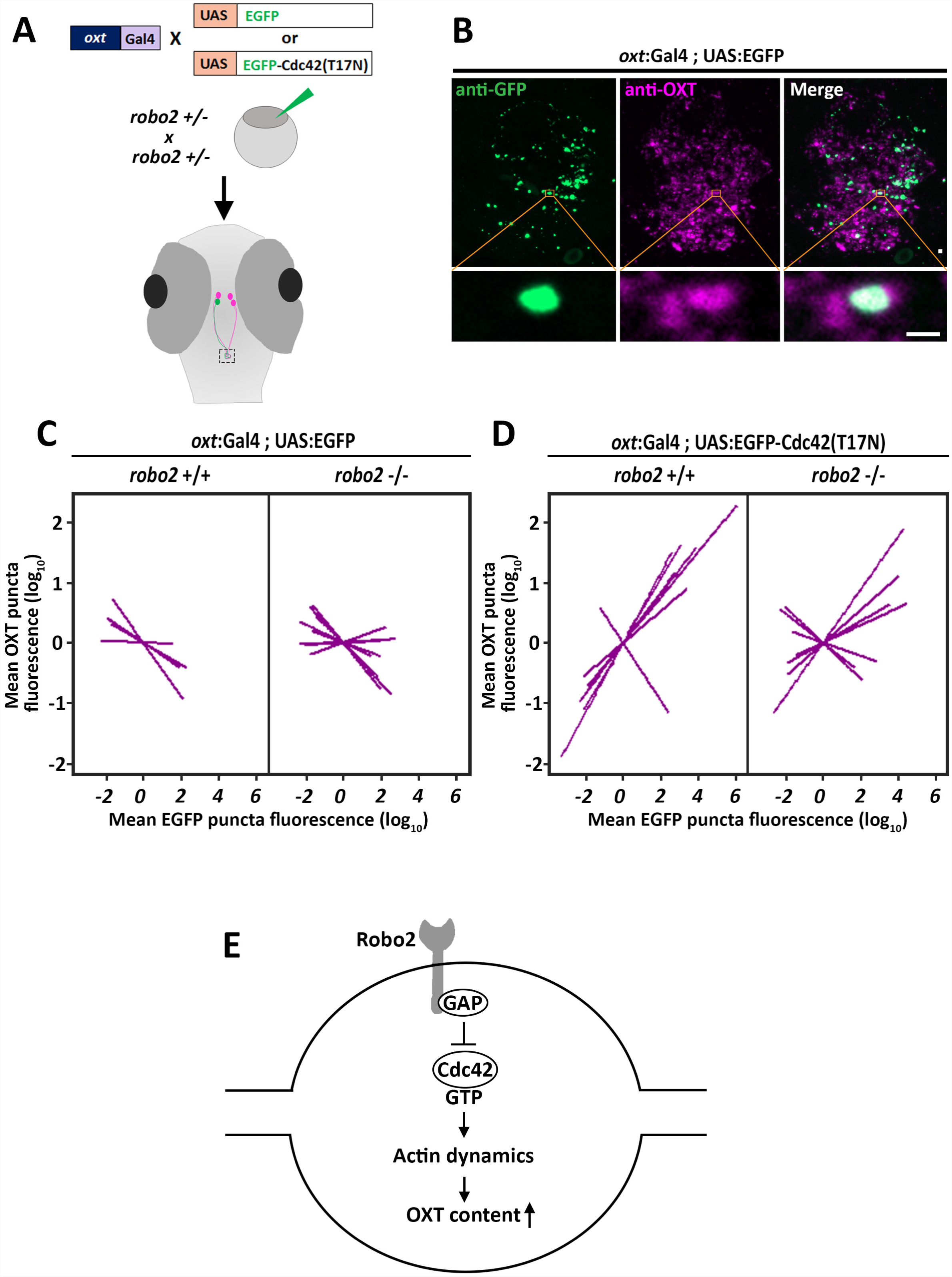
Robo2 regulates synaptic OXT levels via Cdc42. (**A,B**) Assessment of the effect of Oxytocin neuron-specific overexpression of actinregulating protein Cdc42 in Tg(*oxt*:Gal4) larvae. Transgenic embryos expressing the *oxt*:Gal4 driver were injected with transposon-based transgenic vectors containing either control UAS:EGFP or UAS:Cdc42(T17N)-EGFP. Larvae were fixed at 8 days post-fertilization (dpf) and immunostained with anti-GFP and anti-OXT antibodies and neurohypophyseal synapse (NS) were identified as described above. Maximum intensity projection (MIP) reveals mosaic labelling of hypophyseal projecting axons (**B**). Below are magnifications of squared areas showing single stack with colocalization of axonal swelling with OXT puncta. Scale bar 1 µm. (**C,D**) Linear regression analysis comparing between synaptic EGFP levels (mean EGFP puncta) as a function of OXT fluorescence (mean OXT puncta). The data were normalized using mean-centering approach using the scale function in the “R” software. Each line represents a regression line of single animal. The correlation between the mean (log10) fluorescence value of OXT puncta and GFP expression in neurohypophyseal synapse was tested with a linear regression model (ANCOVA), accounting for the effect of GFP fluorescence, and individual fish. Mean EGFP fluorescence is inversely correlated with OXT fluorescence in *robo2+/+* (n=5; p<0.01, adj. R^2^=0.93) and in *robo2-/-*(n=8; and p<0.01, adj. R^2^=0.85) (**C**). Mean EGFP-Cdc42(T17N) fluorescence positively correlates with OXT fluorescence in *robo2+/+* (n=7; p<0.01, adj. R^2^=0.74) larvae but not in in *robo2-/-*(n=7; p=0.35, Adj. R^2^=0.52) (**D**). (**E**) Model of the role of Robo2 in neurohypophyseal synapses: Synaptic Robo signalling inactivates Cdc42 via GAP by affecting the transition of the Rho-GTPase protein Cdc42 from GTPto GDP- bound state. Cdc42 inactivation reduces actin polymerization and increases synaptic OXT content.

## Discussion

To maintain body homeostasis, neurons sense and integrate a multitude of environmental and physiological signals and evoke a response when a deviation is detected. To achieve this, neurosecretory synapses of these cells must maintain adequate levels of neuropeptides that are readily primed to be secreted. This is particularly relevant for neuroendocrine signals that occur at the neurohypophysis level where two neuropeptides that are essential for homeostasis, oxytocin and vasopressin, are released into the blood circulation to exert their effects on peripheral organs (Miyata, 2017; Wircer et al., 2016). Thus, how neuropeptides homeostasis is maintained in such synapses is a fundamental question. In this study, we focused on OXT neuropeptide, which is stored in LDCVs in numerous axonal swellings. These swellings act as *en passant* synapses, which are highly enriched in neurohypophyseal axonal projections (Morris and Pow, 1988). We found that local actin dynamics in those synapses regulate the levels of OXT neuropeptide. We also identified a new signal transduction machinery, Robo2/Cdc42, which regulates synaptic actin state to maintain steady-state OXT content in those synapses.

The majority of reported studies on synaptic OXT content and release were performed on organotypic cultures or by electron microscopy on sliced neurohypophyseal tissues (Alonso et al., 1981; Miyata et al., 2001; Tobin and Ludwig, 2007). Monitoring synaptic OXT content in the transparent zebrafish larvae, we were able to study OXT dynamics in the context of a living vertebrate animal without invasive manipulation. We demonstrated the robustness and validity of our experimental system by showing that synaptic OXT content is depleted upon osmotic challenge, which is in agreement with previous studies done in mammals and also in teleost fish (Alonso et al., 1981; Balment et al., 1980; Neumann et al., 1993; Pierson et al., 1995). Conversely, expression of light chain of botulinum toxin serotype B specifically in the OXT neurons, led to increased synaptic OXT content. This is in agreement with the fact that neurohypophyseal neurons exhibit spontaneous activity and SNARE-mediated synaptic release, which is blocked by the botulinum toxin serotype B (Jurgutis et al., 1996; Tobin et al., 2012).

It was reported that dual pools of actin filaments exist in synapses, a cytoplasmic pool associated with the vesicles and a second pool of cortical filaments associated with the plasma membrane (Alonso et al., 1981; Bleckert et al., 2012). Using super-resolution STORM microscopy, we observed that actin filaments form cage-like structures that surround OXT-containing vesicles. We also observed that actin and OXT exhibit a synaptic Colocalization, possibly forming a readily releasable pool. Our results are in agreement with EM-based studies on rat neurohypophyseal axonal termini (Alonso et al., 1981). As neuropeptides (e.g. OXT) synapses lack active zone, which is typical in small transmitters synapses, the actin cage may regulate vesicle movement and exocytosis. Indeed, synaptic OXT content was decreased upon treatment of zebrafish larvae with cytochalasin D, a cell permeable mycotoxin known to inhibit F-actin formation (Cooper, 1987; Goddette and Frieden, 1986; Lin et al., 1980).

Synaptic actin filaments are dynamic structures undergoing treadmilling and regulating synaptic morphogenesis (Bosch and Hayashi, 2012; Honkura et al., 2008). Actin dynamics was shown to play a role in vesicle clustering in cerebral cortex synapses (Wolf et al., 2015). Using FRAP analysis of the F-actin probe Lifeact-GFP in OXT synapses of live animals, we found that actin state is perturbed in *robo2* mutants. Thus, *robo2* mutants exhibit reduced mobile Lifeact-GFP fraction and increased stable fraction, which was correlated to reduced OXT content in those synapses. Notably, previous studies have shown that Robo2 signalling regulates actin dynamics during neuronal migration and axonal guidance (Kidd et al., 1998; Slovakova et al., 2012; Wong et al., 2001). However, the changes in neurohypophyseal OXT content, which we observed in *robo2*-/- fish were not accompanied by axon guidance defects as the number of neurohypophyseal projecting axons were unaffected.

In view of our findings regarding the involvement of Robo2 in synaptic actin state and OXT content, we hypothesized that changes in synaptic actin mobility regulate the steady-state accumulation of OXT-containing vesicles at the neurohypophyseal release site. To directly test this hypothesis we established a novel transgenic OXT-EGFP fusion line, allowing real time *in vivo* monitoring of the dynamicity of OXT-loaded vesicles. Notably, transgenic rats harbouring a similar OXT-EGFP fusion have already been reported (Hashimoto et al., 2014); however, the deep anatomical location of the neurohypophysis makes it difficult to study the dynamic subcellular processes *in vivo*. Using our zebrafish transgenic reporter, we revealed that Robo2 regulates the accumulation of transiting OXT-loaded vesicles in the synapses. Whether this phenotype is due to decreased vesicle capture or increased exocytosis needs to be further investigated. Interestingly, the majority of the vesicles were immobile, as only ∼10% of nonbleached fluorescent vesicles exchanged into the photobleached synapses. This finding is in agreement with previous work on LDCV mobility in Drosophila neuroendocrine termini (Bulgari et al., 2014).

To link synaptic actin dynamics to neuropeptide content we targeted the intercellular GTPase Cdc42, which acts downstream of Robo signalling pathway to affect actin dynamics. During neuronal migration, Robo signalling promotes Cdc42-GTP hydrolysis that, in turn, attenuates actin polymerization (Wong et al., 2001). We show here that expression of the dominant-negative (i.e. GDP-bound) Cdc42(T17N) in OXT synapses led to increased OXT levels in WT fish in a dose-dependent manner. However, *robo2* mutant was somewhat refractory to the effect of Cdc42(T17N) on synaptic OXT levels, suggesting that Robo2 acts upstream of Cdc42 in the context of synaptic OXT content. This result is consistent with the notion that lack of Robo2 signalling leads to decreased levels of active GTP-bound Cdc42 (Wong et al., 2001). Our findings regarding the regulation of synaptic OXT content might be relevant to other neuropeptidergic and also endocrine cells. Thus, interaction between Cdc42 and vesicle-associated membrane protein 2 (VAMP2) was previously shown in the case of insulin granules in pancreatic beta cells (Nevin and Thormond, 2015). These authors suggested that a mechanism whereby glucose activates Cdc42 to induce the targeting of intracellular Cdc42-VAMP2-insulin vesicles to SNARE proteins at the plasma membrane. Robo2 signalling-dependent actin polymerization is also involved in insulin secretion (Yang et al., 2013).

## Materials and Methods

### Animals

Zebrafish were raised and bred according to standard protocols. All experimental procedures were approved by the Weizmann Institute’s Institutional Animal Care and Use Committee (IACUC). Animals were genotyped by Sanger sequencing. Transgenic zebrafish lines Tg(*oxt*:EGFP)^wz01^ (Blechman et al., 2011), Tg(*oxt*:gal4)^wz06^ (Anbalagan et al., 2018), Tg(*UAS*:Lifeact-EGFP)^mu271^ (Helker et al., 2013), Tg(*UAS*:BotxLCB-EGFP)^icm21^ (Sternberg et al., 2016), Tg(UAS:NTR-mCherry)^c264^ (Davison et al., 2007) and *robo2*^*ti272z*^ (Fricke et al., 2001) were used in this study. See key resources table for oligonucleotide details.

### Transgenesis experiments

We used the Tol2kit transposon-based transgenic vector system for site-specific recombination-based cloning (Kwan et al., 2007) and generated all plasmid DNA constructs. To generate OXTSP-EGFP-OXT construct, sequences were fused by overlap extension PCR and cloned into middle entry Tol2 plasmid 218. The resulting plasmid was recombined downstream of *oxt* promoter in Tol2 pDEST 395 plasmid. The resulting plasmid was coinjected with Tol2 transposase mRNA and founders were outcrossed to obtain germline transmitting lines Tg(*oxt*:OXTSP-EGFP-OXT; *myl7*:EGFP)^wz14^.

### Salt challenge

8- days post-fertilization (dpf) transgenic Tg(*oxt*:EGFP)^wz01^ larvae were treated with 25% artificial sea water (1.75 g Instant ocean Sea salt in 200 mL Danieau buffer) for a period of 60 min and then washed and returned to Danieau’s medium for additional 60 min. The larvae were incubated at 28°C during entire procedure and larvae were fixed in 4% PFA overnight at 4C prior to immunostaining.

### Actin perturbation

For temporal inhibition of actin filaments, a 100 µM stock of cytochalasin D (Sigma C8273) was prepared in DMSO. Briefly, 15-20 Tg(*oxt*:EGFP) 4-dpf larvae were treated with cytochalasin D (200 nM) for 24 h in a 12-well plate at 28°C. Control embryos were treated with equivalent concentrations of DMSO.

For spatial perturbation of Cdc42 signalling in oxytocin neurons, Tg(*oxt*:gal4) embryos were micro-injected with vector *oxt*:gal4 in combination with 10xUAS:EGFP or 10xUAS:*cdc42(*T17N)-EGFP (Ando et al., 2013). All the plasmids were injected at a concentration of 20 ng/μl each and with transposase mRNA at a concentration of 20 ng/μl (∼500 pl/embryo). Using this method, we were able to attain cell labelling in ∼10% of the surviving embryos. 8-dpf larvae that expressed the additionally expressed EGFP heart marker were sorted and fixed in 4% PFA prior to immunostaining. Due to the mosaic nature of the transgenesis experiments and sparse labelling of OXT neurons projecting to the hypophysis (<5% of mosaic clones), transgenesis experiments were performed separately for UAS:EGFP and UAS:Cdc42(T17N)-EGFP constructs.

### *In situ* Hybridization

RNA *in situ* hybridization was performed as described in (Machluf and Levkowitz, 2011). For probe synthesis, partial coding sequences and 3′ UTR of the genes were amplified by PCR, along with a T7 tail in the reverse primer, and purified with PCR cleanup kit. The purified products served as a template to synthesize digoxigenin-labelled antisense mRNA probes using DIG RNA labelling mix (Roche #11277073910).

### Immunofluorescent staining

For immunofluorescent staining, PFA-fixed larvae were washed in PBS (2×10 min), dehydrated using methanol series (25-50-75-100%) and stored at −20°C overnight. The samples were rehydrated from methanol (75-50-25%) to PBS, washed in PBS-Tx (Triton X100, 0.3%; 2×10 min) and blocked in 500 µL of blocking solution (PBS + 10% goat serum + 1% DMSO + 0.3% Triton X100) for 30 min at room temperature. The solution was then replaced with 200 µL of fresh blocking solution with primary antibodies at 1:200 concentration and incubated overnight at 4°C. Samples were washed with PBS-Tx (3×30 min) and treated with 200 µL of secondary antibodies in blocking solution at 1:200 concentration, overnight at 4°C. Then, samples were washed with PBS-Tx (3×30 min), transferred to 75% glycerol (25-50-75%) and the jaws were removed before mounting the larvae ventrally. Rabbit anti-EGFP (ThermoFisher A11122), Guinea Pig anti-OXT (Peninsula labs T-5021) and secondary antibodies were obtained from Jackson ImmunoResearch Laboratories (West Grove, PA).

### Confocal imaging and image analysis

Samples were imaged by using Zeiss LSM 710 or LSM800 confocal microscopes with oil immersion 40X objective. Maximum intensity projection (MIP) images of the whole Z-stacks or subset of Z-stacks were generated using the Zen software (Zeiss). Processing of multiple channel images (i.e., linear adjustments of brightness, contrast and levels) was performed on individual channels using Photoshop CS7 Extended (Adobe). Images were analysed using the open source Fiji image-processing package and Volocity (Perkin Elmer). The number of hypophyseal synapses and OXT puncta was quantified using object measurement tool in Volocity (PerkinElmer) object identifier tool (thresholding was based on SD of fluorescence >=4; size of 0.2 to 25μm^3^). The data were extracted and analysed using R (Team, 2013).

### Fluorescence recovery after photobleaching (FRAP) and image analysis

For live imaging of synaptic Lifeact-EGFP dynamics, 6-dpf live embryos were mounted dorsally in 0.1% low-melt agarose in a 12-mL plate and immersed in Danieau buffer with 0.3% tricaine to prevent movement of the larvae during imaging. The larvae were let to acclimatize for 30 minutes prior to imaging. FRAP experiments were performed using Zeiss LSM 2MP multiphoton microscope with 20X water objective of 1.0 numerical aperture and Chameleon Ti-Sapphire laser (Coherent). The acquisition region was 116 x 116 pixels (28.34 µm^2^), and interval between scanning was 0.35 s at a pixel dwell of 1.76 µs. Four circular ROI of 10 x 10 pixels encircling the synapses were selected for each larva, 3 of them for bleach and 1 for control. Another ROI of similar size was chosen outside the synapses for background quantification. 20 prebleach images were taken and bleach was performed at 75% laser power for 15 iterations and 200 postbleach images were taken.

For live imaging of synaptic OXT-EGFP, embryos were mounted and imaged as described above except for time-lapse parameters. Laser was used at 940 nm 2.5%. The acquisition region was 128 x 64 pixels (14.17 µm x 7.08 µm) and interval between scanning was 1 s at a pixel dwell of 3.15 µs. Two circular ROI of 10 x 10 pixels encircling the synapses were selected for each larva, 1 for bleach and 1 for control. Another ROI of similar size was chosen outside the synapses for background quantification. 20 prebleach images were taken and bleach was performed at 75% laser power and 300 postbleach images were taken. Fluorescence images were drift corrected in Image/Fiji using TurboReg plugin (Thevenaz et al., 1998).

The FRAP values were analysed using EasyFRAP software with full scale normalization to account of difference in synaptic OXT-EGFP fluorescence (Rapsomaniki et al., 2012).

### Dual color three-dimensional STORM Imaging

For STORM imaging, 5-dpf Tg(*oxt*:EGFP) or Tg(*oxt*:OXTSP-EGFP-OXT) embryos were immunostained with anti-OXT (1:200) or anti-EGFP (1:200) as described above. For visualization of actin and OXT, Tg(*oxt*:gal4; UAS:Lifeact-EGFP) larvae were fixed and stained with primary antibodies anti-Neurophysin (PS45) and anti-EGFP at 1:200. Secondary antibody (Alexa 647 or 568, Invitrogen) was used at a concentration at 1:2000. Samples were mounted ventrally without the jaws, and soaked in imaging buffer (50 mM 2-mercaptoethanol, 50 mM Tris-HCl (pH 8.0), 10 mM NaCl, 10% (w/v) glucose), 1x Gloxy for 15 min prior to imaging. 50X Gloxy buffer was made by making 8440 AU glucose oxidase (Sigma #G2133, 50 KU) and 70200 AU catalase (Sigma C40, 100 mg) in 50 mM Tris (pH 8.0) and 10 mM NaCl. Samples were imaged using Vutara SR-200 super-resolution microscope (Bruker). AlexaFluor647 was excited with 640 nm laser (power range of 4-9 kW/cm^2^), AlexaFluor568 was excited with 561 nm laser (6 kW/cm^2^) and 405-nm activation laser power was ramped slowly to maintain optimal singlemolecule density. Images were recorded using a 60x, NA 1.2 water immersion objective (Olympus) and Evolve 512 EMCCD camera (Photometrics) with gain set at 50, frame rate at 50 Hz. Total number of frames acquired was 6000 per labeling dye. Data were analyzed by the Vutara SRX software.

### Western blot

To validate the expression of OXT-EGFP fusion, adult pituitary of transgenic reporter Tg(*oxt*:SP-EGFP-OXT) were dissected and protein extracts were isolated. As controls, Tg(*oxt*:EGFP) pituitary extracts were isolated. Western blotting was performed on isolated protein extracts using anti-EGFP antibodies at 1:1000.

### Statistical analysis

Custom-written R codes were used for analysis of axonal swelling and OXT puncta data. For volume analysis, the values were log-transformed and the mean values were calculated for individual fish. Statistical test were performed between the mean volumes between different genotypes or perturbations. Normality tests were performed and Students’ t-test or Mann-Whitney test or ANOVA were performed.

The correlation between the mean log-transformed (log_10_) fluorescence value of OXT puncta and EGFP axonal swelling was tested with a linear regression model (ANCOVA), accounting for the effect of EGFP axonal swelling, and individual fish.

## Authors’ contributions

S.A., J.B., and G.L. designed the study. S.A. performed immunohistochemistry, confocal imaging and live imaging, quantification and data analysis. J.B. generated transgenic constructs, transgenic lines and performed *in situ* hybridization and biochemical experiments. S.A., and M.G. performed quantification of neurohypophyseal synapses and OXT puncta. S.A. and T.D. performed STORM imaging. R.R assisted in statistical analysis. S.A. and G.L. wrote the manuscript. All authors reviewed the manuscript.

## Acknowledgments

We thank Roy Hofi for animal care; Einav Wircer and Preethi Rajamannar for assisting in transgenic experiments; Joshua Bonkowsky (Univ. of Utah, USA), Noaki Mochizuki (National Cerebral and Cardiovascular Center, Osaka, Japan), Claire Wyart (ICM Institute, Paris) for sharing the *robo2* mutant, Cdc42-related reagents and transgenic UAS:BoTxLCB-EGFP fish respectively. Nitzan Konstantin for English editing. S.A. was supported by Israel PBC-VATAT Fellowship and by Koshland Foundation. G.L. is supported by the Israel Science Foundation (#1511/16), F.I.R.S.T. (Bikura) Individual Grant (#2137/16), Minerva-Weizmann program and the Adelis Metabolic Research Fund. G.L. is an incumbent of the Elias Sourasky Professorial Chair.

---

**Figure 1 Supplement video 1**

Confocal Z-stack images of 5-days post-fertilization (dpf) transgenic reporter zebrafish Tg(*oxt*:EGFP) following immunostaining with anti-EGFP and a specific antibody against endogenous OXT protein.

**Figure 2 Supplement video 1**

Three-dimensional render showing STORM images of endogenous OXT and Lifeact-GFP in neurohypophyseal synapses.

**Figure 4 Supplement video 1**

Three-dimensional render showing STORM images of endogenous OXT in neurohypophyseal synapses of 5-days post-fertilization (dpf) old zebrafish larvae.

## Key resources table

Anbalagan *et al.* Robo2 regulates synaptic oxytocin content by affecting actin state

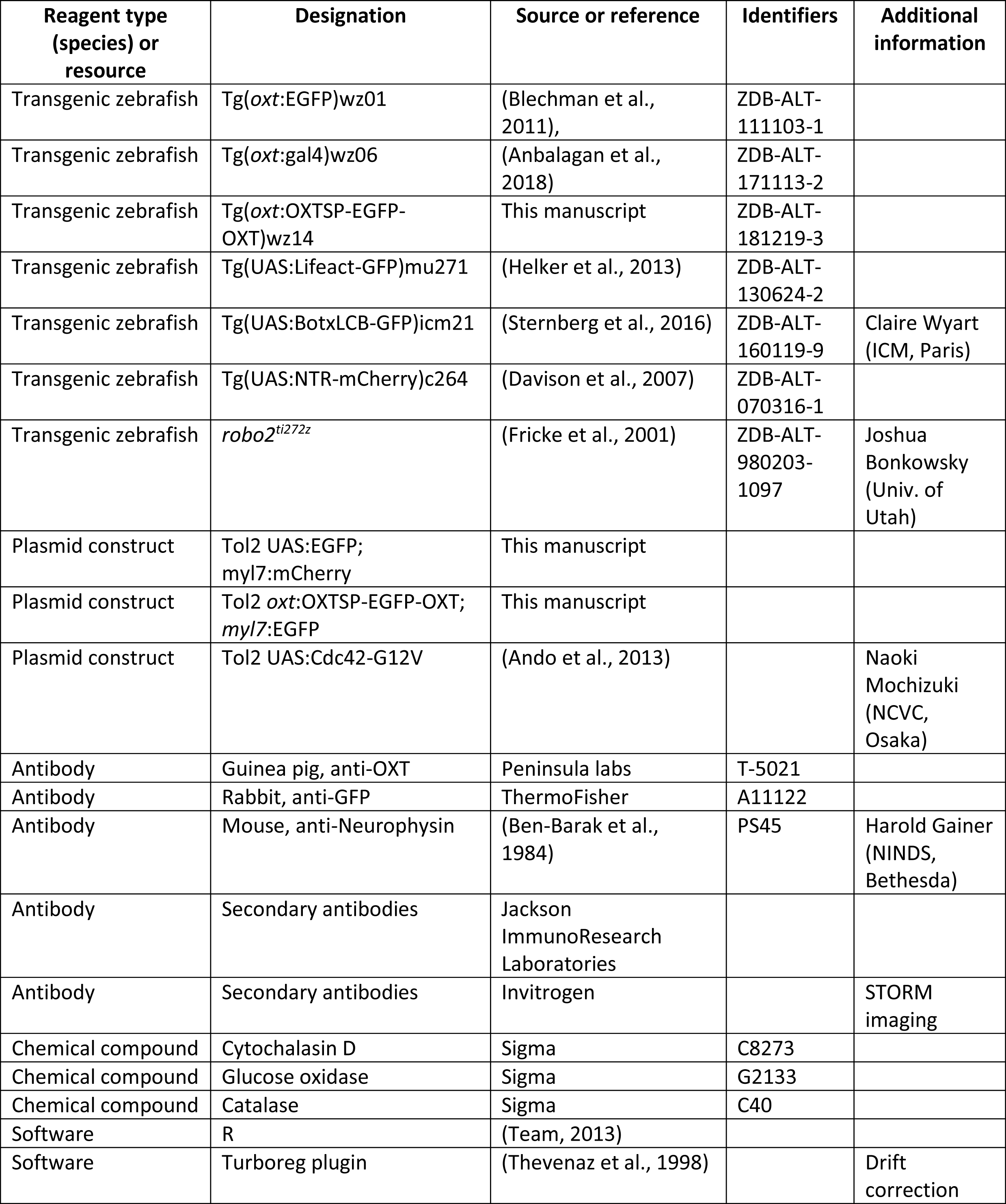
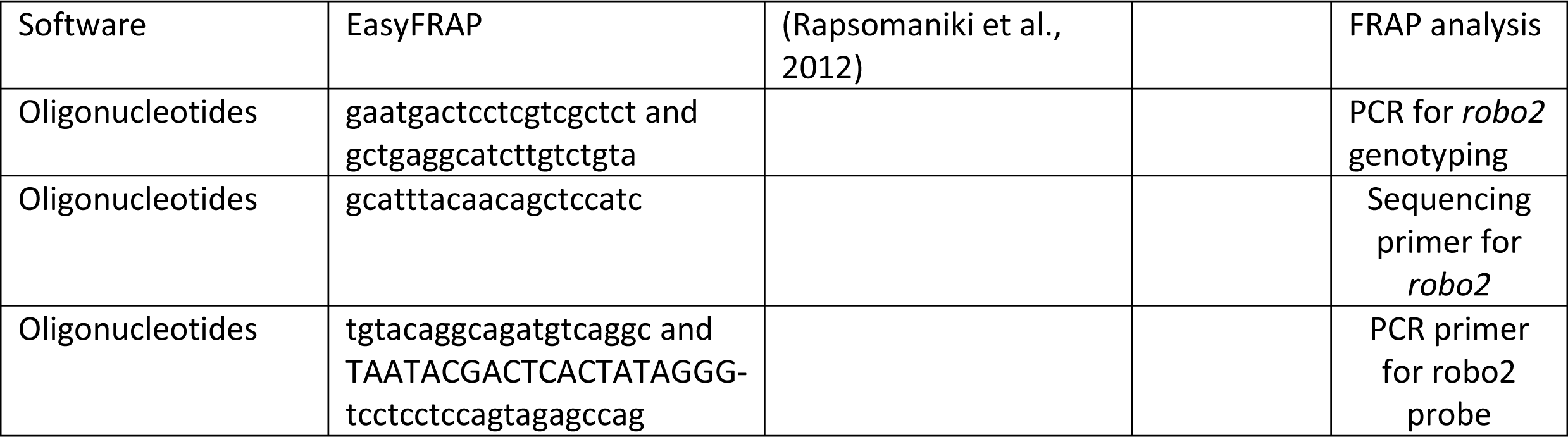

